# The association of longitudinal diet and waist-to-hip ratio from midlife to old age with hippocampus connectivity and memory in old age: a cohort study

**DOI:** 10.1101/2023.12.12.570778

**Authors:** Daria EA Jensen, Klaus P. Ebmeier, Tasnime Akbaraly, Michelle G Jansen, Archana Singh-Manoux, Mika Kivimäki, Enikő Zsoldos, Miriam C Klein-Flügge, Sana Suri

**Author notes:** Corresponding author, Daria EA Jensen.

## Abstract

Epidemiological studies suggest lifestyle factors may reduce the risk of dementia. However, few studies have examined the association of diet and waist-to-hip ratio with hippocampus connectivity. In the Whitehall II Imaging Sub-study, we examined longitudinal changes in diet quality in 512 participants and waist-to-hip ratio in 665 participants. Diet quality was measured using the Alternative Health Eating Index-2010 assessed three times across 11 years between ages 48 and 60 years, and waist-to-hip ratio five times over 21 years between ages 48 and 68 years. Brain imaging and cognitive tests were performed at age 70±5 years. We measured white matter using diffusion tensor imaging and hippocampal functional connectivity using resting-state functional magnetic resonance imaging. In addition to associations of diet and waist-to-hip ratio with brain imaging measures, we tested whether associations between diet, waist-to-hip ratio and cognitive performance were mediated by brain connectivity. We found better diet quality in midlife and improvements in diet over mid-to-late life were associated with higher hippocampal functional connectivity to the occipital lobe and cerebellum, and better white matter integrity as measured by higher fractional anisotropy and lower diffusivity. Higher waist-to-hip ratio in midlife was associated with higher mean and radial diffusivity and lower fractional anisotropy in several tracts including the inferior longitudinal fasciculus and cingulum. Associations between midlife waist-to-hip ratio, working memory and executive function were partially mediated by radial diffusivity. All associations were independent of age, sex, education, and physical activity. Our findings highlight the importance of maintaining a good diet and a healthy waist-to-hip ratio in midlife to maintain brain health in later life. Future interventional studies for the improvement of dietary and metabolic health should target midlife for the prevention of cognitive decline in old age.

## INTRODUCTION

The global trend towards unhealthy dietary habits is leading to an increase in the prevalence of cardiovascular disease, diabetes ^1^, and obesity ^2^, all of which are known risk factors for dementia ^3^. The World Health Organisation (WHO) guidelines recommend a balanced diet with a high plant intake (a Mediterranean-like diet) and weight management to reduce the risk of dementia ^4^. It is therefore important to consider the effect of diet and central obesity on memory and cognition, and associated brain regions such as the hippocampus.

Our recent systematic review revealed several gaps in the literature on the associations of diet and obesity with brain health ^5^. Firstly, most studies have examined single dietary components, but the role of overall diet remains unclear. For instance, studies on older adults have suggested a potentially favourable effect on brain functional connectivity for beetroot juice intake and higher levels of lycopene, ω-3, and ω-6 fatty acids ^5^. Interventions with a caloric-restricted diet ^6^ and resveratrol supplementation^7^ have been associated with higher functional connectivity between the hippocampal subfields and parietal areas.^5^ Similar links between single dietary components and white matter (WM) connectivity have been examined in other studies ^8,9^, but findings are inconsistent. Research on single food components is easier to design and control, but the transferability of conclusions to complete diets is questionable. In the current study, we use comprehensive dietary questionnaires and measured of waist-to-hip ratio (WHR) assessed several times over a 21-year period to address this limitation.

Secondly, although the hippocampus is a key area affected in Alzheimer’s disease, little is known about how it is affected by risk factors such as diet and obesity. While the hippocampus has predominantly been assessed in the context of memory, it might also interact with dietary regulation. Decreased hippocampal activity has been linked to decreased memory of meals (i.e., forgetting when the last meal was consumed), reduced meal intervals and an increased response to food cues ^12^. This can, in turn, lead to increased food intake, poorer dietary quality, and higher body mass index (BMI) ^13^. Moreover, forgetting to eat and the decreased ability to recognise food and eat independently is common in the late stages of Alzheimer’s disease ^14^. While some studies have linked midlife obesity and a poor diet with reduced hippocampal volume, their effects on hippocampal connectivity remain unclear. Here, we use multi-modal brain MRI to examine hippocampal functional and WM connectivity as our primary outcome.

Thirdly, the timing of the diet-brain associations is poorly defined due to the lack of longitudinal studies across the lifespan. Many studies have focussed on either middle-aged or older people, but not on both age groups together ^5^. This is a limitation because the transition period from mid-to-old age (40-70 years old) has been identified as a key window for preventative interventions for dementia risk reduction ^3^.

In this study, we investigated how longitudinal changes in diet quality and WHR during this critical age range are associated with (1) structural and functional connectivity of the hippocampus and (2) cognitive function in later life. Our hypothesis was that higher diet quality and healthier WHR throughout midlife will be related to more favourable indicators of hippocampal functional and structural connectivity measured using resting-state functional MRI (rsfMRI) and diffusion tensor imaging (DTI). We also hypothesised that associations of diet and WHR with cognitive function will be mediated by these brain measures.

## METHODS

### Sample Selection

The Whitehall II (WHII) study, established in 1985 by University College London, is a longitudinal study of 10,308 individuals from the British Civil Service who have been followed up for over 30 years through 13 study waves ^21^. WHR was measured at waves 3 (1991-1994), 5 (1997-1999), 7 (2002-2004), 9 (2007-2009) and 11 (2012-2013), while diet was assessed at waves 3, 5, and 7. A subset of participants also received brain MRI scans and cognitive tests as part of the WHII Imaging Sub-study conducted at the University of Oxford in 2012-2016 ^22^. MRI scans were obtained shortly after wave 11 for 775 participants according to the published protocol ^22^. For this study, we included participants of the Imaging Sub-study who had information on diet from at least one previous wave, WHR from at least two previous waves and good quality structural, rsfMRI and DTI scans (i.e., no excessive head motion artefacts or blurry images, or incidental findings such as large strokes, tumours, or brain cysts; see appendix p2). The final sample for the fMRI analysis consisted of 665 and 512 participants for analyses of WHR and diet respectively For the DTI analyses, 8 (WHR) and 6 (diet) participants were additionally excluded because of missing or poor-quality DTI data, resulting in a final sample of 657 and 506 participants for WHR-DTI and diet-DTI analyses respectively.

Informed written consent was obtained from all participants at each data collection of the Whitehall II study in procedures approved by the University of Oxford Central University Research Ethics Committee (Application reference: MS IDREC-C1-2011-71) and the University College London Medical School Committee on the Ethics of Human Research (reference: 85/0938). Ethical approval was obtained in accordance with the Declaration of Helsinki (1975, revised in 1983).

### AHEI-2010 score

Participants filled in a Food Frequency Questionnaire (FFQ) with 240 questions, which assessed 127 food items consumed in an average week. This self-report questionnaire assessed items such as how much fruit, vegetables, and drinks they had consumed on a scale from 1-5 within the last seven days. The FFQ was administered repeatedly over 30 years in waves 3, 5, 7, and 9 of the study, however, due to excessive missing information at Wave 9, we only used data from waves 3, 5, and 7 for this study. Diet quality was measured using the AHEI-2010 score ^20,23^ (for details see appendix p 2).

### Waist-to-hip ratio

While most studies have used BMI as a proxy marker for obesity, more recent literature has highlighted WHR as a potentially more accurate predictor of health and disease ^24^. Here, WHR was estimated by dividing the waist circumference at the biggest diameter by the trochanter circumference. We used WHR data that were obtained before brain scans, including waves 3, 5, 7, 9 and 11.

### Longitudinal changes in diet and waist-to-hip ratio

We used linear mixed effect models with maximum likelihood estimation (LME function, *NLME* package in R version 4.2.1) to extract the intercept (projected diet at baseline), the slope (linear change in diet over time), and the quadratic slope (non-linear change in diet over time) in diet across three waves and 11-year follow-up. The same method was used to assess change in WHR across five waves and a 21-year follow-up. Both the intercepts and slope (time) were fitted as random effects (see model equations in appendix p 2). We allowed for correlations between time points, allowing a continuous autoregressive process for a continuous time covariate. We tested whether the addition of a quadratic term for time improved model fit. The best-fit model was the linear model for diet, whereas the quadratic term improved significantly the model fit for WHR (see appendix p 3).

### MRI analysis

The 775 participants of the WHII Imaging Sub-study were scanned on a 3T scanner at the Wellcome Centre for Integrative Neuroimaging (FMRIB Centre, Oxford). Of them, 552 participants were scanned on a 3T Siemens Magnetom Verio scanner (Erlangen, Germany) with a 32-channel head coil (April 2012 - December 2014). After a scanner upgrade, the remaining 223 were acquired using a 3T Siemens Prisma Scanner (Erlangen, Germany) with a 64-channel head–neck coil in the same centre (July 2015-December 2016; detailed protocol in ^22^). The scan parameters were identical or closely matched between scanners (for MRI details, see appendix pp 4-5), and a scanner model covariate was used in all analyses.

WM microstructure was assessed using DTI scans analysed with tract-based spatial statistics (*FSL-TBSS)*^25^. Global fractional anisotropy (FA), radial diffusivity (RD), axial diffusivity (AD), and mean diffusivity (MD) were extracted from the mean TBSS skeleton (details see appendix p 4). We additionally extracted FA, MD, RD, and AD values from three regions of interest in proximity to the hippocampus based on the literature ^26^: the fornix, the inferior longitudinal fasciculus (ILF), and the cingulum.

Hippocampal functional connectivity was analysed using seed-based correlation analyses with a hippocampus kernel sphere of 4mm radius (detailed description in appendix pp 4-5 *randomise*, ^27^) with 1000 permutations. The group-level voxel-wise analysis included strict threshold-free cluster enhancement (*TFCE*) and a correction for family-wise errors (*FWE*, p<0·05) for multiple voxel-wise comparisons. Mean hippocampal connectivity values were extracted from significant clusters and plotted for visualisation.

### Cognitive tests

Cognitive tests were performed at the time of the MRI scan. We examined three cognitive domains: working memory, fluency, and executive function. The working memory domain included the verbal episodic short-term memory recall of words on the Hopkins Verbal Learning Test-Revised (HVLT-R) and the total score on the digit span forward, backward, and sequence tests (DST). Fluency included semantic fluency: number of animals named in one minute; lexical fluency: number of words beginning with the letter F in one minute; executive function included the difference in time to completion between Trail Making Test B and A and the total score on the digit coding test (DCOD) adapted from the Wechsler Adult Intelligence Scale-Fourth Edition (WAIS-IV). Overall cognitive health was determined using the Montreal Cognitive Assessment (MoCA).

### Statistical analysis

To examine the association between the intercepts and slopes of AHEI-2010/WHR with (1) structural connectivity, (2) functional connectivity, and (3) cognitive outcomes we performed linear regression models using R (*stats* package in R with function *lm*).

We performed a causal mediation analysis using the *mediation* package in R to test whether the association of AHEI-2010/WHR on cognitive function (working memory, fluency, and executive function) at the MRI Phase was mediated by MRI markers (rsfMRI and DTI). We only tested the mediation when individual paths between the intercepts and slopes of AHEI-2010/WHR with DTI and/or rsfMRI measures were identified as statistically significantly associated and further significant correlations with cognitive outcome measures were observed (see further details in appendix p 5). Analyses were corrected for multiple comparisons using the Bonferroni correction. Statistical significance was accepted at p<0·017 for the WM region of interest analysis across three pre-defined tracts, and p<0·0083 for the six cognitive tests.

All analyses were adjusted for sex, MRI scanner model, age, years of education, BMI, mean arterial pressure, physical activity, and the Montreal Cognitive Assessment (MoCA) score all measured at the MRI timepoint. RsfMRI analyses were corrected for head motion and voxel-wise grey matter density confounder. AHEI-2010 analyses we additionally adjusted for total energy intake (kcal/day; further details see appendix p 5).

All analyses were performed using R, version 2.3.1 and FSL, version 6.0.5.

## RESULTS

Participants’ characteristics for the sample used in the functional connectivity analyses are shown in Table 1. Mean outcome measures (extracted WM metrics and cognitive performance) are in appendix pp5-6 and mean hippocampus connectivity maps are shown in appendix p 6.

**Table 1.** *Characteristics of the participants*^*\**^.

|  | Wave 3<br>1991-1994 | Wave 5<br>1997-1999 | Wave 7<br>2002-2004 | Wave 9<br>2007-2009 | Wave 11<br>2011-2013 | MRI scan<br>2012-2016 |
| --- | --- | --- | --- | --- | --- | --- |
| **AHEI-2010 sample included in functional connectivity analyses** |  |  |  |  |  |  |
| N (% female) | 512 (21.29%) |  |  |  |  |  |
| Age (years) | 47.79 ± 5.19 | 53.57 ± 5.16 | 59.03 ± 5.14 | .. | .. | 69.81 ± 5.11 |
| AHEI-2010 score (all participants) | 55.61 ± 9.44 | 56.19 ± 9.66 | 56.25 ± 10.27 | .. | .. | .. |
| (women) | 56.73 ± 9.43 | 56.75 ± 9.93 | 56.98 ± 10.32 |  |  |  |
| (men) | 55.24 ± 9.41 | 56.02 ± 9.66 | 56.01 ± 10.20 |  |  |  |
| Education (years) | .. | .. | .. | .. | .. | 14.01 ± 3.08 |
| Physical activity (met/week) | .. | .. | .. | .. | .. | 2748.46 ± 1822.31 |
| Mean energy intake | 2202.59 ± 504.43<br>(Average across waves 3,5, and 7) |  |  |  |  | .. |
| Mean arterial pressure | .. | .. | .. | .. | .. | 98.88 ± 12.05 |
| BMI | .. | .. | .. | .. | .. | 26.24 ± 4.13 |
| MoCA | .. | .. | .. | .. | .. | 27.15 ± 2.40 |
| **WHR sample included in functional connectivity analyses** |  |  |  |  |  |  |
| N (% female) | 664 (19.88%) |  |  |  |  |  |
| Age (years) | 47.71 ± 5.14 | 53.47 ± 5.12 | 58.96 ± 5.09 | 63.88 ± 5.10 | 67.94 ± 5.10 | 69.75 ± 5.07 |
| WHR (all participants) | 0.89 ± 0.08 | 0.91 ± 0.07 | 0.93 ± 0.08 | 0.93 ± 0.07 | 0.95 ± 0.07 | .. |
| (women) | 0.89 ± 0.08 | 0.91 ± 0.07 | 0.93 ± 0.08 | 0.94 ± 0.07 | 0.95 ± 0.07 |  |
| (men) | 0.89 ± 0.08 | 0.91 ± 0.07 | 0.93 ± 0.08 | 0.93 ± 0.08 | 0.95 ± 0.07 |  |
| Education (years) | .. | .. | .. | .. | .. | 14.11 ± 3.11 |
| Physical activity (met/week) | .. | .. | .. | .. | .. | 2750.70 ± 1810.21 |
| Mean arterial pressure | .. | .. | .. | .. | .. | 98.75 ± 11.83 |
| BMI | .. | .. | .. | .. | .. | 26.07 ± 4.14 |
| MoCA | .. | .. | .. | .. | .. | 27.23 ± 2.28 |

Abbreviations: AHEI-2010 - Alternative-Health Eating Index 2010, WHR - waist-to-hip ratio, met- Metabolic Equivalent of Task, MoCA - Montreal Cognitive Assessment.
\*Measures are specified as their mean ± their standard deviation. Diet was measured using the AHEI-2010 score across three waves (3,5, and 7) and an MRI scan was acquired in N=512 participants 11 years later. Waist-to-hip ratio (WHR) was measured across 21-years in five waves (3, 5, 7, 9, and 11) and the MRI scan was acquired in N=664 participants.

The AHEI-2010 score increased linearly over the 11-year period (Fig 1A, β_1_=0·06 (95%CI=[-0·02, 0·13]), p=0·15). The increase was small and not statistically significant, indicating minimal improvement in diet quality during midlife. By contrast, WHR increased non-linearly and statistically significantly over the 21-year period (Fig 1B, β_1_=0·0046 (95%CI=[0·0038, 0·0054], p<0.001, β_2_=- 0·0001 (95%CI=[-0·0001, -0·0000], p<0·001).

**Figure 1:**
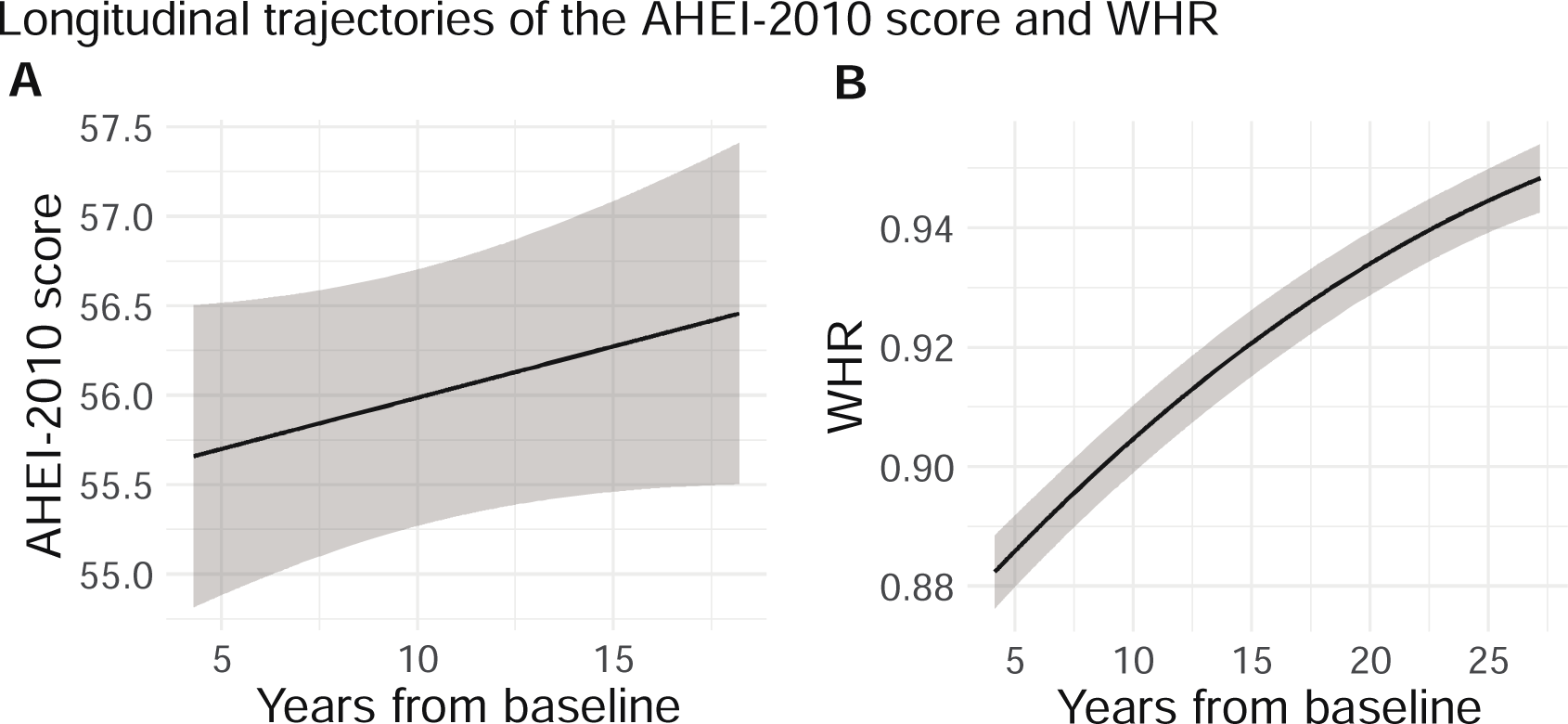
Predicted longitudinal trajectories of the AHEI-2010 score (N=506) and WHR (N=664). **A** shows the linear increase of the AHEI-2010 score across 11 years (Waves 3, 5, and 7) with a slope of β_1_=0·06 (95% CI=[-0·02, 0·13]) and a baseline level of β_0_=55·41, (95% CI=[54·35, 56·47]). **B** shows the quadratic increase of WHR across 21 years (Waves 3, 5, 7, 9, and 11) with a slope of β_1_=0·0046 (95% CI=[0·004-0·01]), a baseline level of β_0_=0·86 (95% CI=[0·86, 0·87]), and a quadratic term of β_2_=-0·0001 (95% CI=[-0·0001, -0·0000]). Abbreviations: AHEI-2010 - Alternative-Health Eating Index 2010, WHR - waist-to-hip ratio.

Voxel-wise analyses showed that a higher increase in AHEI-2010 across 11 years (i.e., higher slope) was associated with higher FA, lower MD, and lower AD in several WM tracts (Table 2, Fig 2). Higher FA was observed in the corticospinal tract, superior thalamic radiation, frontal aslant tract, and frontal regions. A less pronounced association with MD was observed in the optic radiation and the superior parietal lobe. A very localised association with AD was found in the superior longitudinal fasciculus (Fig 2).

**Table 2:** Results of the voxel-wise association of AHEI-2010 and WHR with FA, MD, RD, and AD.*

| Outcome | Cluster size<br>(mm<sup>3</sup>) | Cluster<br>number | Number of<br>voxels | p-value at<br>maxima | Coordinates<br>of maxima (x<br>y z) | Location of maxima |
| --- | --- | --- | --- | --- | --- | --- |
| AHEI-2010 (slope) |  |  |  |  |  |  |
| FA | 13424 | 1 | 891 | 0.027 | 28 -31 32 | R Corticospinal Tract |
|  |  | 2 | 640 | 0.03 | -23 -9 35 | L Superior Thalamic<br>Radiation |
|  |  | 3 | 126 | 0.043 | 27 24 26 | R Frontal Aslant Tract |
|  |  | 4 | 12 | 0.048 | 32 32 29 | R Middle Frontal Gyrus |
|  |  | 5 | 9 | 0.046 | 31 33 36 | R Frontal Lobe |
| MD | 296 | 1 | 25 | 0.043 | 18 -77 9 | R Optic Radiation |
|  |  | 2 | 7 | 0.038 | -61 -24 11 | L Temporal Lobe |
|  |  | 3 | 5 | 0.044 | 18 -73 43 | R Superior Parietal Lobe |
| AD | 2272 | 1 | 282 | 0.036 | -34 -44 31 | L Superior Longitudinal<br>Fasciculus |
|  |  | 2 | 2 | 0.048 | 18 -73 43 | R Superior Parietal Lobe |
| WHR (intercept) |  |  |  |  |  |  |
| MD | 317656 | 1 | 39689 | 0.001 | 6 24 13 | Corpus Callosum Body |
|  |  | 2 | 12 | 0.047 | 50 -16 45 | Postcentral Gyrus |
|  |  | 3 | 6 | 0.047 | 31 -41 -5 | Corpus Callosum Body |
| RD | 239120 | 1 | 29890 | 0.001 | -8 24 -6 | Genu of Corpus Callosum |

Abbreviations: AHEI-2010 - Alternative-Health Eating Index 2010, WHR - waist-to-hip ratio, AD - axial diffusivity, FA - fractional anisotropy, MD - mean diffusivity, RD - radial diffusivity, R - right, L - left.
\*Cluster sizes (total size in mm<sup>3</sup> and the number of significant voxels), threshold-free cluster enhancement (TFCE) and family-wise error (FWE) corrected *p*-values, voxel coordinates in MNI-152 space, and locations of cluster maxima are reported for all significant clusters.

**Figure 2:**
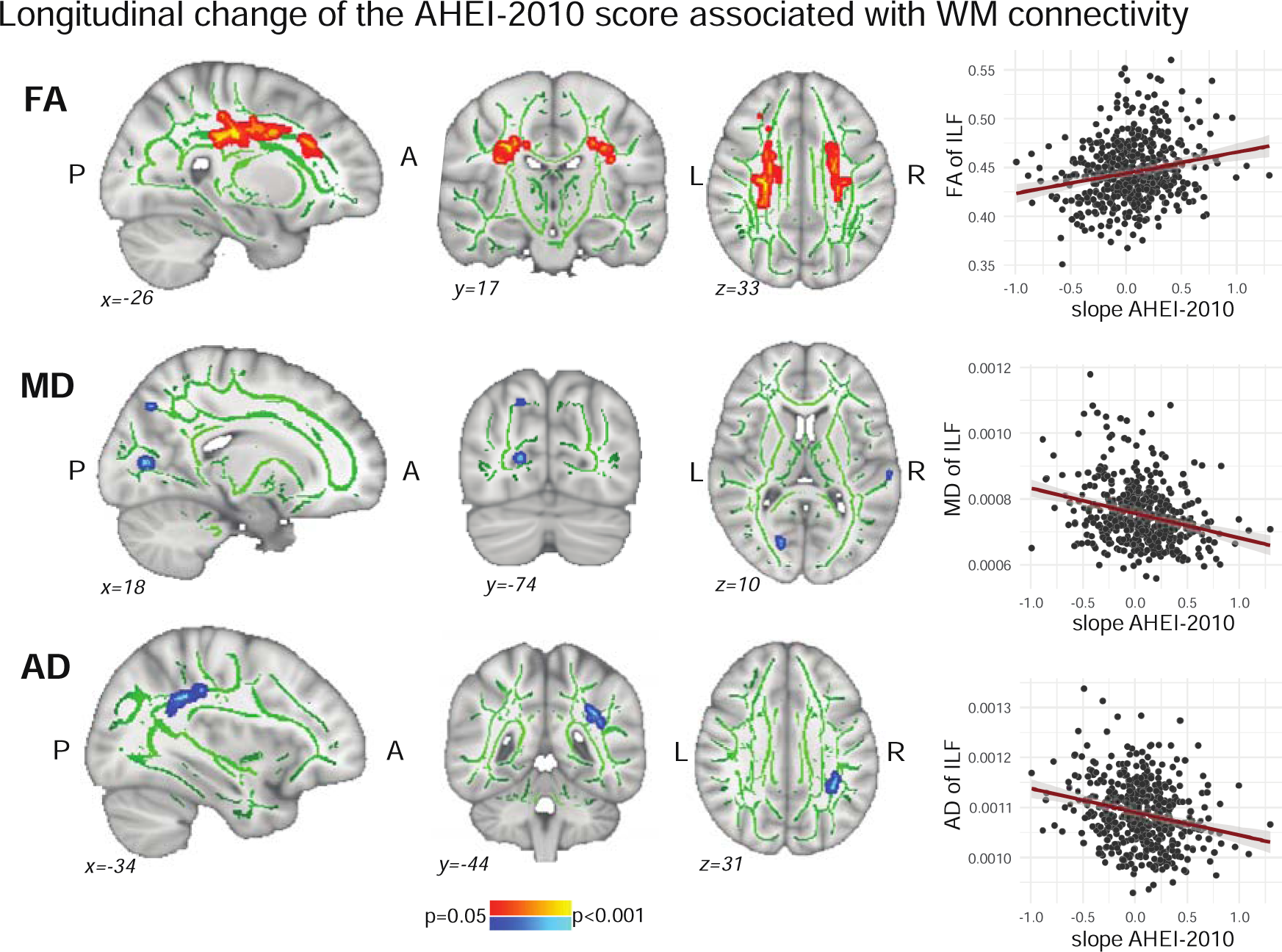
Voxel-wise association between the slope of the AHEI-2010 score and WM connectivity of **A** fractional anisotropy (FA) **B** mean diffusivity (MD) and **C** axial diffusivity (AD). The left panels show FWE-corrected TFCE statistical maps overlaid on the FMRIB58_FA standard image. Green tracts represent the mean FA skeleton. Associations of the higher AHEI-2010 slopes were positive (in red) with FA and negative (in blue) with MD and AD. The right panels show scatter plots (for visualisation purposes only) of extracted WM measures (FA, MD, or AD) from the specific cluster correlated with the slope of AHEI-2010. Abbreviations: AHEI-2010 - Alternative-Health Eating Index 2010, S - superior, I - inferior, L - left, R - right, A - anterior, P – posterior, WM - white matter.

We found that higher WHR in midlife (i.e., higher intercept) was associated with higher MD and RD covering 28·81% and 21·69% of the total WM tracts, respectively (Fig 3A). The maxima were identified in the corpus callosum and the postcentral gyrus (Table 2).

**Figure 3:**
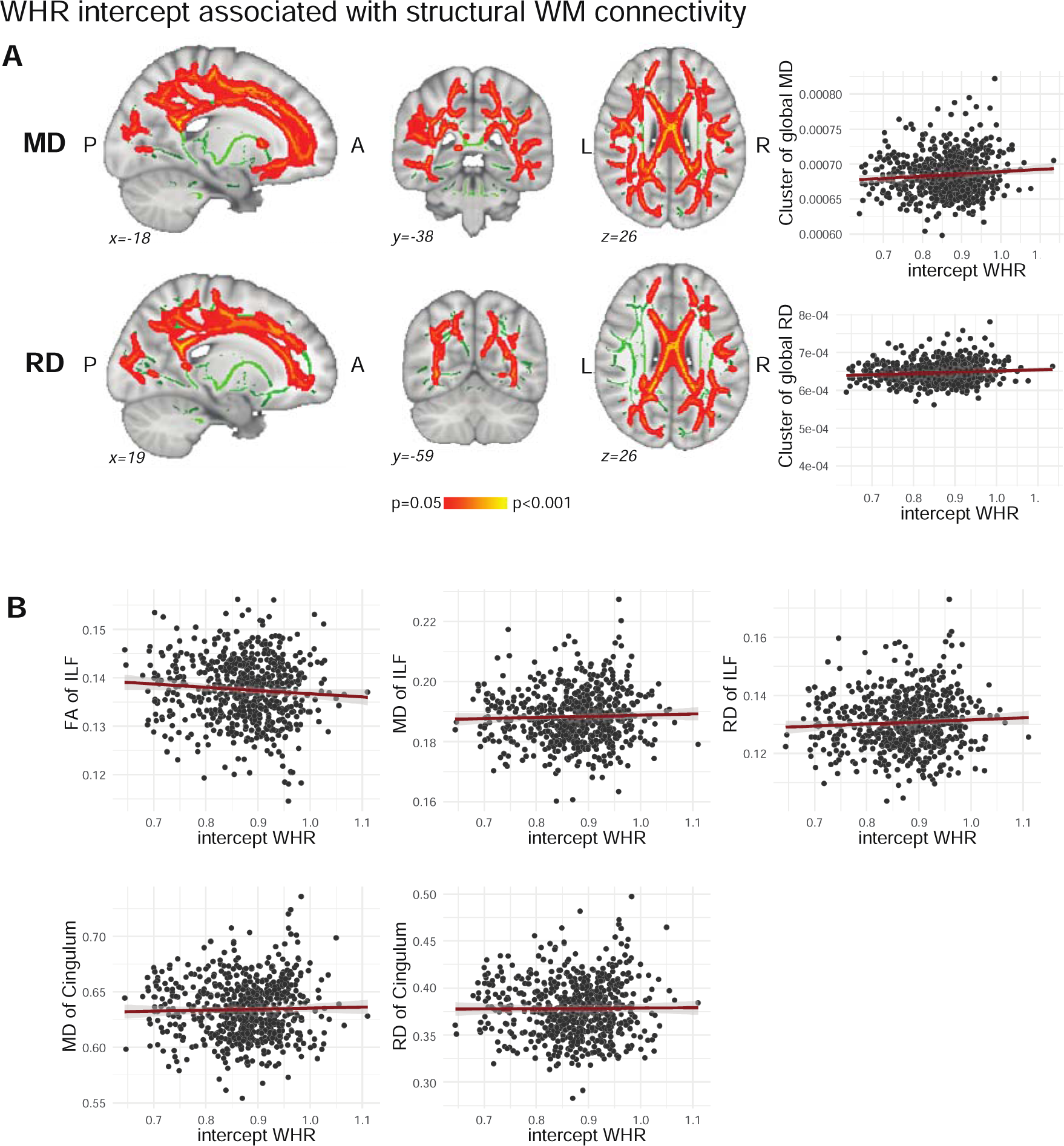
A Voxel-wise association between the intercept of WHR and WM connectivity of mean diffusivity (MD) and radial diffusivity (RD). The left panels show FWE-corrected TFCE statistical maps overlaid on the FMRIB58_FA standard image. Green tracts represent the standardized mean FA skeleton. Associations between the increase of WHR across time were positive (in red) with MD and RD. The right panels show scatter plots (for visualisation purposes only) of extracted WM measures (MD or RD) from the specific cluster correlated with the intercept of WHR. **B** A separate ROI analysis with three pre-defined hippocampal WM tracts showed associations between the intercept of WHR with ILF and Cingulum connectivity. The partial regression plots represent significant association which survived Bonferroni correction for multiple comparisons (p<0·017). Abbreviations: WHR - waist-to-hip ratio, S - superior, I - inferior, L - left, R - right, A - anterior, P – posterior, WM - white matter.

The three pre-defined hippocampal tracts were also significantly associated with WHR intercepts and AHEI-2010 slopes (Table 3). However, only associations between higher WHR intercepts and lower FA, higher MD, and RD in the inferior longitudinal fasciculus, as well as higher MD and RD in the cingulum survived the Bonferroni correction for multiple comparisons across the three tracts (p<0·017, Fig 3B).

**Table 3:** Results of the region of interest analysis showing associations of the fornix, inferior longitudinal fasciculus, and cingulum with WHR intercept and AHEI-2010 slope. P-values are markers in bold for p<.05 and with an ^*^ for p<.017 for the Bonferroni correction of multiple comparisons across the three white matter tracts.

|  | WM tract | WM | Beta (SE) | CI | p-value |
| --- | --- | --- | --- | --- | --- |
| **WHR intercept** | Fornix | FA | -0.06 (0.03) | -0.11, -0.01 | **0.026** |
|  |  | MD | 0.02 (0.01) | -0.00, 0.04 | 0.057 |
|  |  | RD | 0.01 (0.01) | -0.00, 0.03 | 0.056 |
|  |  | AD | 0.03 (0.01) | -0.00, 0.05 | 0.075 |
|  | ILF | FA | -1.26 (0.35) | -1.95, -0.56 | **0.00042\*** |
|  |  | MD | 0.82 (0.26) | 0.31, 1.34 | **0.0018\*** |
|  |  | RD | 0.90 (0.25) | 0.42, 1.38 | **0.0003\*** |
|  |  | AD | 0.37 (0.23) | -0.07, 0.81 | 0.10 |
|  | Cingulum | FA | -0.20 (0.09) | -0.37, -0.02 | **0.029** |
|  |  | MD | 0.29 (0.10) | 0.09, 0.48 | **0.0042\*** |
|  |  | RD | 0.24 (0.09) | 0.07, 0.40 | **0.0057\*** |
|  |  | AD | 0.09 (0.06) | -0.04, 0.21 | 0.18 |
| **AHEI-2010 slope** | Fornix | FA | -0.38 (0.19) | -0.74, -0.01 | **0.043** |
|  |  | MD | 0.16 (0.07) | 0.03, 0.30 | **0.020** |
|  |  | RD | 0.13 (0.06) | 0.02, 0.25 | **0.022** |
|  |  | AD | 0.25 (0.11) | 0.04, 0.46 | **0.018** |
|  | ILF | FA | 4.25 (2.32) | -0.31, 8.81 | 0.068 |
|  |  | MD | -4.00 (1.70) | -7.34, -0.65 | **0.019** |
|  |  | RD | -3.60 (1.62) | -6.78, -0.42 | **0.027** |
|  |  | AD | -3.06 (1.48) | -5.98, -0.14 | **0.040** |
|  | Cingulum | FA | 0.83 (0.58) | -0.32, 1.97 | 0.16 |
|  |  | MD | -1.32 (0.67) | -2.64, 0.01 | **0.049** |
|  |  | RD | -1.08 (0.56) | -2.18, 0.02 | 0.06 |
|  |  | AD | -0.38 (0.43) | -1.23, 0.47 | 0.38 |

Abbreviations: ILF - inferior longitudinal fasciculus, WHR - waist-to-hip ratio, AHEI-2010 - Alternative-Health Eating Index 2010, WM - white matter, AD - axial diffusivity, FA - fractional anisotropy, MD - mean diffusivity, RD - radial diffusivity.

No associations with WM were observed for the intercept of AHEI-2010 or the quadratic slope of WHR.

We observed significant associations of hippocampal functional connectivity with diet but not WHR. Higher AHEI-2010 intercepts (i.e., better projected midlife diet quality) were associated with higher functional connectivity between the left hippocampus and occipital lobe and cerebellum, and one small cluster between the right hippocampus and cerebellum (Table 4, Fig 4).

**Table 4:**
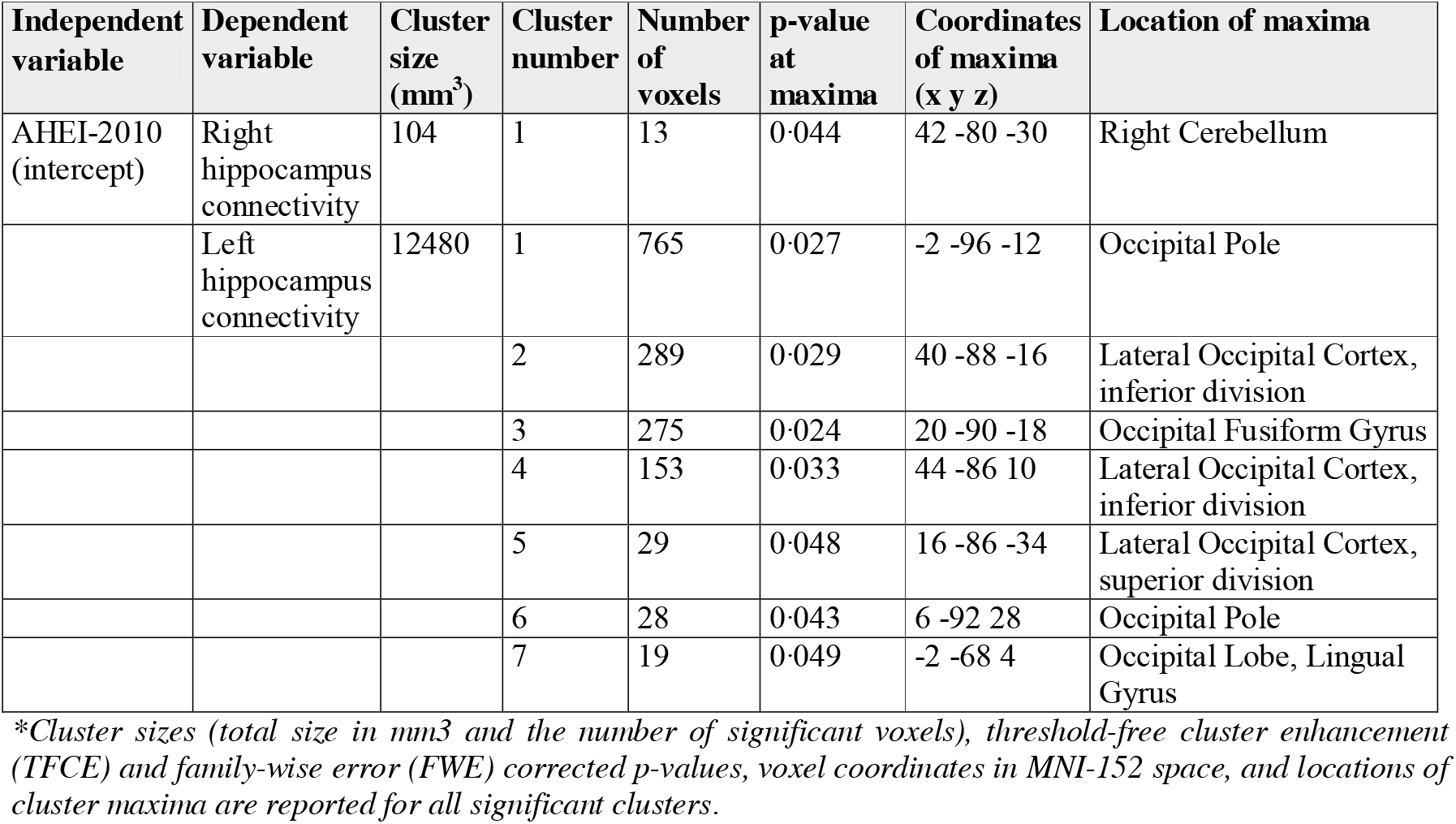
Cluster report for seed-based correlation analysis. The Table shows associations between the intercept of the AHEI-2010 score with functional connectivity of the right and left hippocampus*.

| Independent variable | Dependent variable | Cluster size (mm<sup>3</sup>) | Cluster number | Number of voxels | p-value at maxima | Coordinates of maxima (x y z) | Location of maxima |
| --- | --- | --- | --- | --- | --- | --- | --- |
| AHEI-2010 (intercept) | Right hippocampus connectivity | 104 | 1 | 13 | 0.044 | 42 -80 -30 | Right Cerebellum |
|  | Left hippocampus connectivity | 12480 | 1 | 765 | 0.027 | -2 -96 -12 | Occipital Pole |
|  |  |  | 2 | 289 | 0.029 | 40 -88 -16 | Lateral Occipital Cortex, inferior division |
|  |  |  | 3 | 275 | 0.024 | 20 -90 -18 | Occipital Fusiform Gyrus |
|  |  |  | 4 | 153 | 0.033 | 44 -86 10 | Lateral Occipital Cortex, inferior division |
|  |  |  | 5 | 29 | 0.048 | 16 -86 -34 | Lateral Occipital Cortex, superior division |
|  |  |  | 6 | 28 | 0.043 | 6 -92 28 | Occipital Pole |
|  |  |  | 7 | 19 | 0.049 | -2 -68 4 | Occipital Lobe, Lingual Gyrus |

\*Cluster sizes (total size in mm<sup>3</sup> and the number of significant voxels), threshold-free cluster enhancement (TFCE) and family-wise error (FWE) corrected p-values, voxel coordinates in MNI-152 space, and locations of cluster maxima are reported for all significant clusters.

**Figure 4:**
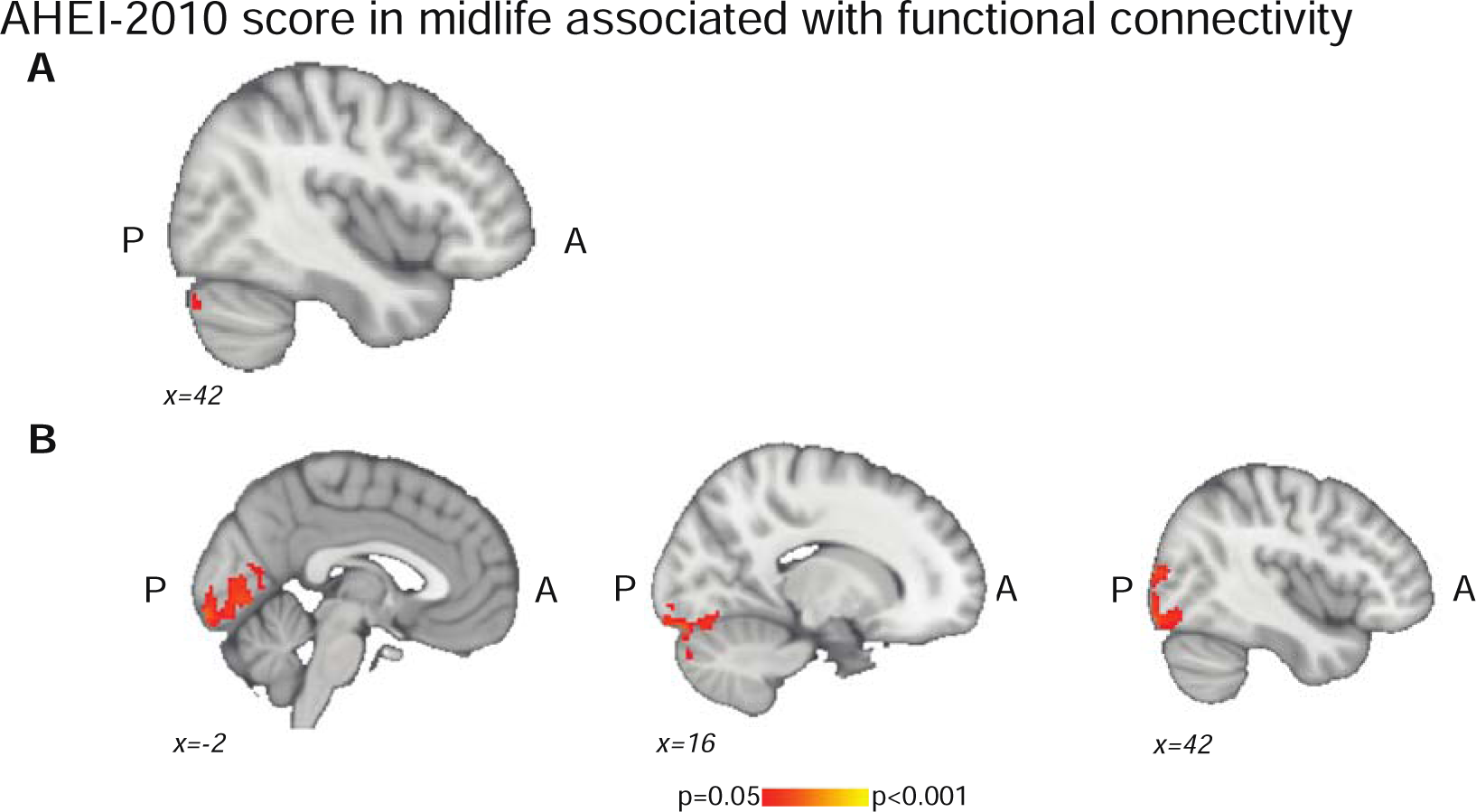
The association between the intercept of the AHEI-2010 score and hippocampal functional connectivity in N=512 participants. **A** Higher functional connectivity of the right hippocampus to a small cluster in the right cerebellum was associated with higher AHEI-2010 scores in midlife (21 years before the MRI scan). **B** Higher functional connectivity of the right hippocampus to bigger clusters in the occipital lobe and the right cerebellum were associated with higher AHEI-2010 scores in midlife. Images show FWE-corrected TFCE statistical maps overlaid on the MNI-152 template. Abbreviations: AHEI-2010 - Alternative-Health Eating Index 2010, A - anterior, P - posterior.

Higher WHR intercept was significantly associated with lower cognitive performance on the verbal episodic memory (β=-0·0014, p=0·0048), digit span (β=-0·0013, p=0·00079), and digit coding test (β=-0·0006, p=0·00071, Fig 5A), after correcting for confounders. The associations survived a Bonferroni-corrected threshold for significance (p<.0083, correcting across six tests; appendix pp 6-7).

**Figure 5:**
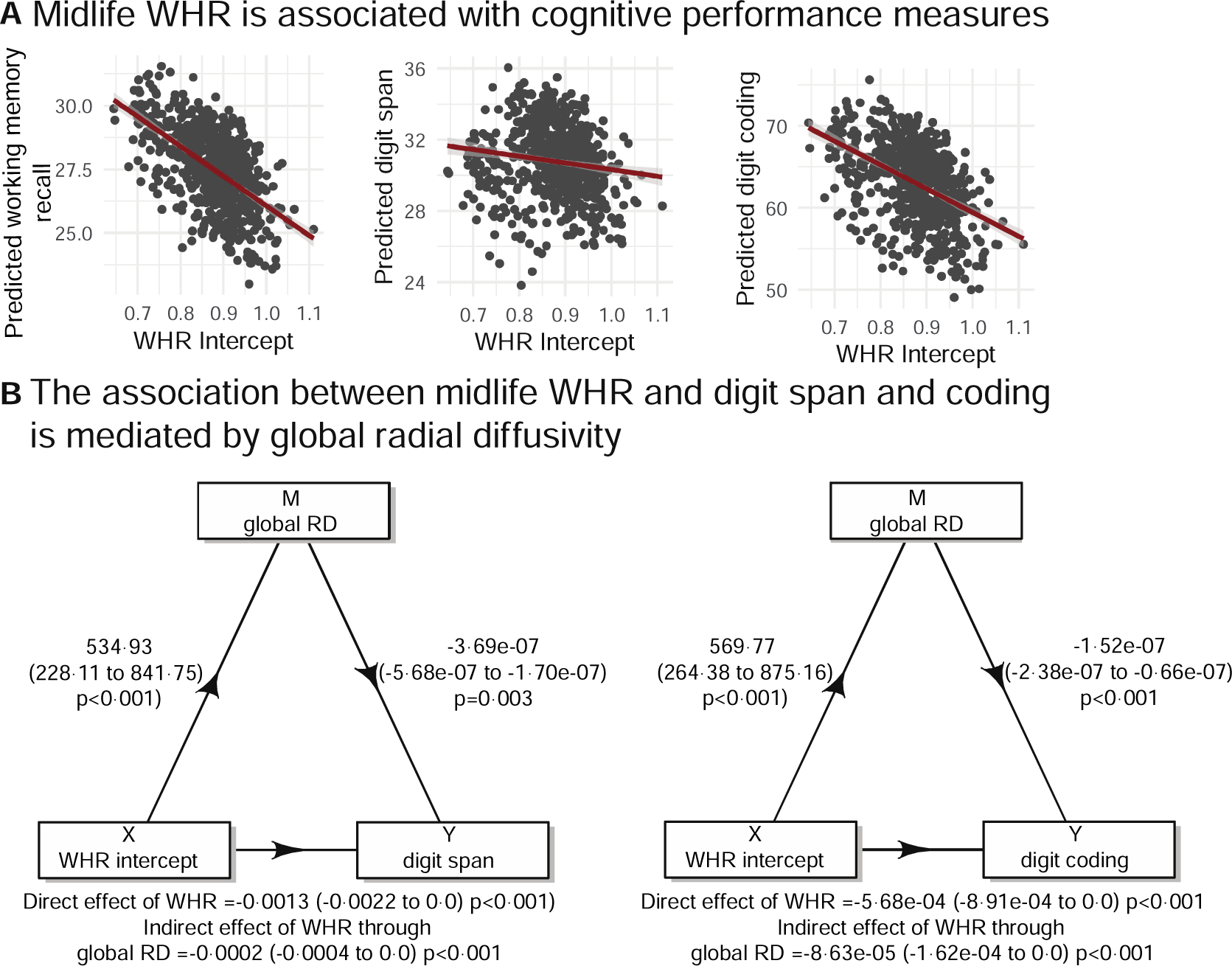
Association between WHR and cognitive performance, mediated by white matter integrity. **A** Partial regression plots show the association between the intercept of WHR and the predicted cognitive outcomes, which survived Bonferroni correction for multiple comparisons (p<0·017). Higher WHR in midlife was associated with lower cognitive performance on verbal episodic memory (β=-0·0014, p=0·0048), digit span (β=-0·0013, p=0·0008), and digit coding tests (β=-0·0006, p=0·0007). **B** Causal mediation analysis showed that the WHR in midlife (i.e. intercept) has a direct and indirect effect on digit span and digit coding, which was partially mediated by global white matter RD. Abbreviations: WHR - waist-to-hip ratio, RD - radial diffusivity.

We then tested whether these associations were mediated by the MRI variables found to be associated with WHR intercepts (see Table 3). Therefore, tested mediators were WM connectivity measures: global (MD and RD), ILF (FA, MD, and RD), and cingulum (MD and RD). We found that the association of WHR intercepts with digit span and digit coding was mediated via the global RD (digit span: b=-0·0002, 95% CI=[-0·0004, 0·0], p<0·001; digit coding: b=-00009, 95% CI=[-0·0002, 0·0], p<0·001; see Fig 5B; appendix pp 6-7). Specifically, 15% (95% CI=[0·04, 0·38], p<0·001 and 95% CI=[0·05, 0·43], p<0·001) of the effect of WHR in midlife on the performance in digit span and digit coding tests respectively was mediated via higher global RD. Other significant mediation effects did not survive the Bonferroni correction for multiple comparisons for seven tested mediators (i.e., WM connectivity measures; p<0·0071, see appendix pp 7-8).

Additionally, the WHR slope was associated with digit span (β=0·0001, p=0·023), and the AHEI-2010 slope was associated with lexical fluency (β=0·0073, p=0·024), but neither of these associations survived the Bonferroni threshold (p<0.0083, appendix pp 6-7). The AHEI-2010 intercept was not associated with any cognitive domain, hence no mediations were performed.

## DISCUSSION

Our findings show that better diet quality and healthier WHR in midlife and improved trajectories throughout mid-to-old age were linked to stronger structural and functional connectivity of the hippocampus in older age. Furthermore, lower WHR in midlife was associated with better working memory and executive function later in life, and this pathway was mediated by WM diffusivity. Overall, these findings may have implications for optimizing the timing of dietary and metabolic interventions aimed at maintaining brain and cognitive health during the lifespan.

Diet quality showed a non-significant increase during the 11-year follow-up period and the average AHEI-2010 score did not meet the cut-off of a ‘healthy’ diet at any Wave. The mean AHEI-2010 score level at baseline was 55·61 out of the optimal dietary score of 110 (i.e., <80% of the optimal score). The guidelines for similar dietary scores such as the Health Eating Index 2010 ^28^ have shown that on a scale from 0-100, a score below 51 (below 50% of the optimal dietary score) indicates an overall “poor” diet quality, whereas scores above 80 indicate a “good” healthy diet. Thus, our score at baseline indicated generally unhealthy diets amongst the participants. Previous studies in the Whitehall II cohort ^20^, have suggested that the lower AHEI-2010 scores may be influenced by higher alcohol consumption in this cohort. The modest improvement in the AHEI-2010 score over time could be influenced by access to healthier foods or improved socio-economic status in older age or retirement.

While previous studies have found positive associations between high diet quality and larger hippocampal volume ^17,20^, this is to our knowledge the first longitudinal study showing that better diet quality in midlife is associated with higher hippocampal functional connectivity in older age. We noted higher hippocampal connectivity to the occipital lobe and cerebellum. While these are distant connections of the hippocampus, their volumes have previously been associated with diet ^30–33^. However, we note that these associations although significant, were small and very localised, and hence warrant replication and cautious interpretation.

We found that the (modest) improvements in the AHEI-2010 score (i.e., higher slopes) across an 11-year period from mid-to-old age was associated with better WM integrity (higher FA, and lower MD and AD). We identified higher FA in widespread tracts (e.g., corticospinal tract, superior thalamic radiation), lower MD in the optic radiation and the superior parietal lobe and lower AD in the superior longitudinal fasciculus (SLF). These regions have previously been implicated as markers for WM microstructural damage in ageing and dementia ^34^. Our findings are also in line with studies showing that higher ω-3 fatty acid levels ^8^ and a general healthier diet in older adults are linked to higher FA and lower MD in the SLF and corpus callosum and higher global FA ^5^. Taken together, we suggest that strategies to improve diet quality and adherence to current dietary guidelines in midlife may possibly reduce the risk of related disorders such as dementia.

We observed that WHR generally increased non-linearly during a 21-year period from mid-to-old age in both men and women. WHR higher than 1·0 for men and 0·86 for women is classified as high risk ^35^ of developing cardiovascular disease. In this cohort, we noted that on average WHR increased from 0·89 (mean age 47·7) to 0·95 at the final follow-up (mean age 67·9) in men and women. While previous studies have revealed largely consistent associations between high BMI and low WM integrity from midlife to older age ^5,36,37^, associations with more precise measures of abdominal fat such as WHR have been understudied. Here, we report that higher WHR in midlife predicts widespread increases in diffusivity of the WM, covering up to 22-29% of all WM tracts. Analyses of specific WM tracts linked to the hippocampus revealed lower FA, higher MD, and RD in ILF and higher MD and RD in the cingulum with higher midlife WHR. Our longitudinal findings are in line with previous cross-sectional studies showing associations between higher WHR and lower FA in several WM tracts, including the corpus callosum and ILF in older adults ^36^ and cingulum in middle-aged adults ^37^. This ILF and cingulum are known to be implicated in Alzheimer’s disease ^34^, and our findings suggest that they may also be especially relevant for WHR-related brain health changes.

Our study had several limitations. First, dietary data were collected using a semi-quantitative FFQ, which can be open to self-report errors. Nevertheless, the FFQ covers a wide range of foods and has been validated. Second, diet quality was assessed by AHEI-2010 which might not cover all aspects of a “healthy” diet and may not be adapted to the dietary habits of all populations; however, the AHEI-2010 score has already been established and used in previous studies ^20^.

Our analyses were adjusted for sex, but our study is limited as the cohort is predominately male and we lacked the statistical power to examine women and men separately (<20% women). The literature shows sex differences in individuals’ WHR, body fat and abdominal fat ^35^. One study found that body fat was positively, while abdominal fat was negatively associated with cortical thickness in older men but not women ^38^. Furthermore, we observed that the baseline WHR was above the high-risk threshold for women, but only at moderate risk for men for developing cardiovascular-related diseases ^35^. These sex-specific differences in risk highlight the importance of balanced or stratified studies.

Our cohort is also predominantly White British, well-educated, and generally healthier than the UK population ^39^. This limits the generalisability of our findings, particularly as longitudinal studies can suffer from a survival bias, and since MRI cohorts tend to create a healthier-than-average participant bias due to the selection of participants.

In conclusion, we found that diet and WHR in midlife were associated with later-life hippocampal functional and WM connectivity. We also showed that higher WHR in midlife was associated with working memory and executive function in older age and this association was partially mediated by WM diffusivity. The findings from this longitudinal cohort study suggest that interventions to improve diet and manage central obesity might be best targeted in mid-life (48-70 years old) in order to see beneficial effects on brain and cognitive health in older age.

## Author contributions

Conceptualization: Daria EA Jensen, Sana Suri.

Data curation: Daria EA Jensen, Sana Suri, Enikő Zsoldos.

Formal analysis: Daria EA Jensen, Sana Suri.

Funding acquisition: Klaus P. Ebmeier, Archana Singh-Manoux, Mika Kivimäki.

Investigation: Daria EA Jensen.

Methodology: Daria EA Jensen, Sana Suri, Michelle G Jansen, Klaus P. Ebmeier, Miriam C Klein-Flügge.

Project administration: Enikő Zsoldos, Klaus P. Ebmeier.

Resources: Tasnime Akbaraly, Michelle G Jansen.

Supervision: Sana Suri, Klaus P. Ebmeier, Miriam C Klein-Flügge.

Validation: Daria EA Jensen, Sana Suri.

Visualisation: Daria EA Jensen.

Writing – original draft: Daria EA Jensen, Sana Suri.

Writing – review & editing: Daria EA Jensen, Sana Suri, Klaus P. Ebmeier, Tasnime Akbaraly, Miriam C Klein-Flügge, Archana Singh-Manoux, Mika Kivimäki, Enikő Zsoldos.

## Ethics approval and Consent to participate

Ethical approval was obtained in accordance with the Declaration of Helsinki (1975, revised in 1983). Informed written consent was obtained from all participants at each data collection of the Whitehall II study in procedures approved by the University of Oxford Central University Research Ethics Committee (Application reference: MS IDREC-C1-2011-71) and the University College London Medical School Committee on the Ethics of Human Research (reference: 85/0938).

## Consent to publish

For the purpose of open access, the author has applied a CC BY public copyright license to any Author Accepted Manuscript version arising from this submission. The views expressed are those of the authors and not necessarily those of the NHS, the NIHR or the Department of Health and Social Care.

## Data sharing agreement

All data that support the findings of this study are available by application to the Whitehall II Study and the Whitehall II Imaging Sub-Study on the DPUK portal (https://portal.dementiasplatform.uk/Apply). As restrictions apply to the availability of these data, which were used under license for the current study, the authors cannot publicly share this data.

Code that allows DPUK users to replicate analyses, including plotting all figures presented in the manuscript will be provided on OSF once this manuscript is accepted for publication. Intermediate analysis outputs can be made available to registered DPUK users upon request and after approval of a proposal. Please see the README file in the Scripts folder for further details.

## Acknowledgements

We thank all the participating civil service departments; the British Occupational Health and Safety Agency; the British Council of Civil Service Unions; all participating civil servants in the Whitehall II Study; and all members of the Whitehall II Study team at University College London who so helpfully collaborated with us. The Whitehall II Study team comprises research scientists, statisticians, study coordinators, nurses, data managers, administrative assistants, and data entry staff, who make the study possible. We are grateful to the staff at the Wellcome Centre for Integrative Neuroimaging in Oxford, in particular research radiographers Michael Sanders, MSc, Jon Campbell, MMRTech, BcAppSc, Caroline Young, DCR(R), David Parker, BSc(Hons), who acquired the scans. Martin R. Turner, MA, MBBS, PhD, FRCP (Wellcome Centre for Integrative Neuroimaging, Oxford, United Kingdom), and his colleagues advised on incidental findings and taking over clinical responsibility for such participants. No compensation was provided for staff contributions to this study.

