## Supplementary for "The association of longitudinal diet and waist-to-hip ratio from midlife to old age with hippocampus connectivity and memory in old age: a cohort study"

**Appendix 1: Flowchart of sample selection**

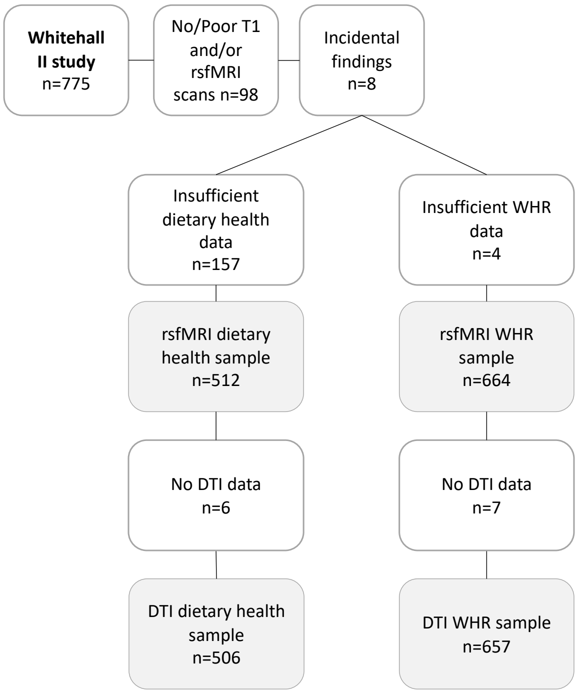

**Appendix 2: Details of the AHEI-2010 score**

The AHEI-2010 score [1] is based on eleven components, where a higher score is related to a higher intake of vegetables, fruit, whole grains, nuts, legumes, long chain ω-3 fats, and polyunsaturated fatty acids, avoidance or low intake of sugar-sweetened drinks and fruit juice, red and processed meat, trans-fat, sodium and avoidance or low consumption of alcohol. Each component was scored from 0 to 10 points, where a score of 10 points indicated that the recommendations were fully met and a score of 0 represented the least healthy dietary behaviour. All the component scores were summed to obtain the total AHEI-2010 score [1].

**Appendix 3: Linear Mixed Effect Model Equations**

As dependent variables in the linear mixed effect model, we used WHR values from five waves (21 years) and the dietary AHEI-2010 score [1] over three waves (11 years). In each linear mixed effect model, we employed the following equations for each dependent variable:

1) Linear Model

$$val_n = \beta_0 + \beta_1 t_n + e_n$$

2) Quadratic Model

$$val_n = \beta_0 + \beta_1 t_n + \beta_2 t_n^2 + e_n$$

$val$ = WHR/diet value is the dependent variable for each participant ($n$) at each timepoint
$\beta_0$ = random intercept
$\beta_1$ = random linear slope
$\beta_2$ = random quadratic slope
$t$ = time to baseline (Wave 3) for each wave, $t^2$ is the squared time variable.
$e$ = is the residual error.

### Appendix 4: Comparison of likelihoods of fitted models

***Table A1***: *Comparison of likelihoods of fitted models (L-linear and Q-quadratic) of WHR and the AHEI-2010 score. In WHR, but not the AHEI-2010 model, the quadratic slope significantly improved the model fit.*

| | Model | df | AIC | BIC | *Log-Likelihood* | Test | Likelihood ratio | p-value |
|---|---|---|---|---|---|---|---|---|
| **WHR** | L | 7 | -11817·70 | -11774·95 | 5915·852 | | | |
| | Q | 11 | -11986·51 | -11919·32 | 6004·254 | 1 vs 2 | 176·8046 | <0·0001 |
| **AHEI-2010** | L | 7 | 10841·04 | 10878·40 | -5413·521 | | | |
| | Q | 11 | 10847·66 | 10906·37 | -5412·832 | 1 vs 2 | 1·379315 | 0·8478 |

*Abbreviations: df - degree of freedom, AIC - Akaike's Information Criteria, BIC - Bayesian Information Criteria, L - linear mixed effect model, Q - quadratic mixed effect model, AHEI-2010 - Alternative Health Eating Index 2010.*

### Appendix 5: MRI image acquisition and pre-processing details

***Table A2***: *Whitehall II cohort details and MRI acquisition parameters.*

| **Cohort** | **Whitehall II cohort (WHII)** | |
|---|---|---|
| **Study design** | Cross-sectional and longitudinal diet/WHR data | |
| **Healthy Participants** | N=775 | |
| **Age range** | 60-85 years (Wave 11) | |
| **Scanner** | 3T - Verio | 3T - Prisma |
| **Structural MRI** | | |
| **T1-sequence** | ME-MPRAGE | MPRAGE |
| **TR (ms)** | 2,530 | 1,900 |
| **TE (ms)** | 1·79/3·65/5·51/7·37 | 3·97 |
| **TI (ms)** | 1,380 | 904 |
| **Flip angle** | 7° | 8° |
| **Voxel dimension (mm$^3$)** | 1x1x1 | 1x1x1 |
| **Field of view (mm)** | 256 | 192 |
| **Acquisition time (min:sec)** | 6:12 | 5:31 |
| **Resting-state fMRI** | | |
| **Sequence** | Multiband | |
| **TR (ms)** | 1300 | |
| **TE (ms)** | 40 | 91 |
| **Flip angle** | 66° | |
| **Voxel dimension (mm3)** | 2x2x2 | |
| **Field of view (mm)** | 212 | |
| **Number of volumes** | 460 | |
| **Acquisition time (min:sec)** | 10:10 | |

*Abbreviations: T - Tesla, TR - repetition time, TE - echo time, TI - inversion time, ME-MPRAGE - multi-echo magnetization-prepared rapid acquisition with gradient echo, WHR - wait-to-hip ratio.*

T1-weighted structural MRI (multi-echo MPRAGE sequence with motion correction, TR=2530ms, TE=1·79/3·65/5·51/7·37ms, flip angle=7°, FOV=256 mm, voxel dimension=1mm isotropic, acquisition time=6min 12s), multiband echo-planar imaging resting-state fMRI scans (voxel=2mm isotropic, TR=1·3s, acquisition time=10min 10s, multi-slice acceleration factor=6, number of volumes=460) and diffusion-weighted images using an echo planar sequence, with 60 diffusion-weighted directions ($b$-value=1500s/mm$^2$), 5 non-diffusion weighted images ($b$-value=0s/mm$^2$) and one B0 volume in the reversed phase-encoded direction (TR=8900ms, TE=91·2ms, FOV=192mm, voxel dimension=2mm isotropic) were analysed for this study.

MRI data was pre-processed using FSL version 6.0.5 [2,3] as described in Filippini & collegues [4]. Bias correction using FSL-ANAT, brain extraction, and partial-volume tissue segmentation using FMRIB Automated Segmentation Tool (*FAST*, [5]) were performed on T1 scans. Resting-state fMRI data were pre-processed using motion correction, brain extraction, high-pass temporal filtering, and field-map correction tools in FEAT. Non-neuronal fluctuations were regressed out of the 'signal' using single-subject independent component analysis (ICA) and automatic component classification using FMRIB's ICA-based X-noiseifier (*FIX*, [6,7]). Diffusion-weighted scans were pre-processed using the FMRIB diffusion toolbox (*FDT*), which included motion correction and correction for eddy currents with *FSL-TOPUP* [8]. Diffusivity and anisotropy maps were extracted using *DTIfit* and aligned into standard space using FMRIB's Nonlinear Registration Tool (*FNIRT*).

### Appendix 6: Further details on the analysis of structural connectivity using DTI and hippocampal functional connectivity using rsfMRI

#### 1. Hippocampal structural connectivity using DTI

WM microstructure was assessed using DTI scans analysed with tract-based spatial statistics (*FSL-TBSS)* [9]. DTI detects the directionality of diffusion of water molecules within the axons. This diffusion is unrestricted along the axon, hindered perpendicularly due to the presence of the myelin sheath. The directionality and diffusivity were qualified by DTI parameters such as fractional anisotropy (FA), radial diffusivity (RD), axial diffusivity (AD), and mean diffusivity (MD). If the diffusion in a voxel is anisotropic this means that it follows more easily along the axons as perpendicular to them. These DTI parameters give indirect indicators of fibre tract integrity and have been found to differentiate between mild cognitive impairment, Alzheimer's disease, and other dementias (see review [10]). Global FA, MD, RD and AD were extracted from the mean TBSS skeleton, with values ranging from 0 (representing isotropic diffusion) to 1 (representing anisotropic diffusion). We additionally extracted FA, MD, RD and AD values from three regions of interest in proximity of the hippocampus based on the literature [11]: the fornix, the inferior longitudinal fasciculus (ILF), and the cingulum.

#### 2. Hippocampal functional connectivity using rsfMRI

Hippocampal functional connectivity was analysed using seed-based correlation analyses. For the right and left hippocampus masks obtained from FIRST [12], we created a binarised seed mask using a kernel sphere with a radius of 4mm, a threshold of 0·75 in structural space (see figure A1). Then, each hippocampus mask was registered from structural to resting-state fMRI space using the FSL *applywarp* command.

For each subject, we used *first-level FEAT* analyses to calculate seed-based connectivity maps of both the right and left hippocampus to the rest of the brain. Within each *first-level FEAT*, a General Linear Model (GLM) was fitted to the data. In each GLM, we used two contrasts: one to determine brain regions positively correlated with the seed and one for negative correlations. As we were interested in the connectivity of hippocampus seed with grey matter, we regressed out the signal from white matter and cerebrospinal fluid (CSF) using white matter and the CSF time series as confounders in the *first-level FEAT* for each subject. This produced a hippocampal partial correlation (or functional connectivity) map for each subject, which shows regions that are significantly positively and negatively correlated with the left and right hippocampus.

To identify between-subject differences, individual functional connectivity maps were used for the *higher-level FEAT* analysis. We created GLMs for (1) the intercept, (2) the linear slope (for WHR and AHEI-2010), and (3) the quadratic slope (for WHR only). Between-subject differences were investigated using nonparametric permutation inference (FSL's tool *randomise,* [13]) with 1000 permutations. Analyses were restricted to voxels within the grey matter (using the MNI152 template thresholded to 0·3).

A 4D covariate image was generated to obtain grey matter density masks using the command *feat_gm_prepare* in FSL. This grey matter density image was used as a voxel-dependent confounder (in addition to all other confounders listed in ED8). The group-level voxel-wise analysis included strict threshold-free cluster enhancement (*TFCE*) and a correction for family-wise errors (*FWE*, p<0·05) for multiple voxel-wise comparisons. Mean hippocampal connectivity values were extracted from significant clusters and plotted for visualisation.

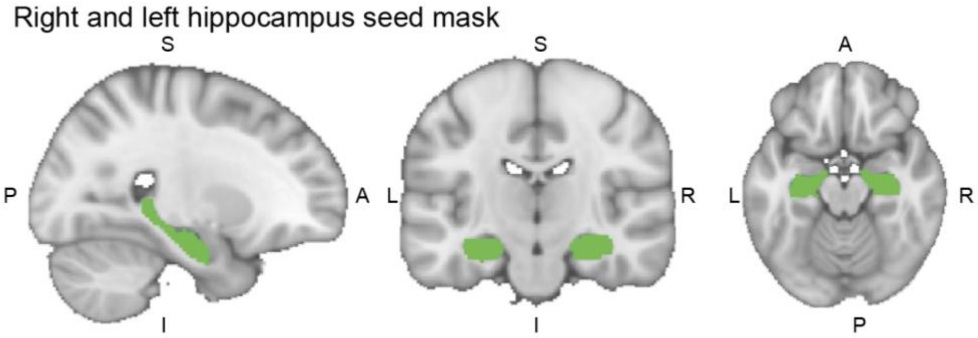

***Figure A1***: *Hippocampal seed mask used for the seed-based correlation analysis in example subject. MNI coordinates in this view: x=33, y=54, z=27.*

**Appendix 7: Mediation Analysis**

We performed a causal mediation analysis using the *mediation* package in R to test whether the association of WHR (intercept and slope across 21 years) and/or dietary quality (i.e., AHEI-2010 intercept and slope across 11 years) on cognitive performance (in six tests) at the MRI Phase was mediated by MRI markers (rsfMRI and DTI).

This was used to test the statistical significance of the direct effect from the indirect (i.e., mediated) effect of diet/WHR on cognitive performance. Thus, diet/WHR intercept/slope was set as the X-variable, MRI variables were set as mediator (M) variables, and the cognitive performance measures were set as outcome variables (Y).

The mediation analysis was performed on participants with complete data only when there was a significant association between X and Y. We ran two separate multiple linear models to assess the individual path of the indirect effect. The first model had cognitive performance as the dependent variable (Y), and the brain MRI variable as the mediator variable (M), and other covariates (see Methods) as independent variables (IVs; Path: M + covariates → Y). In the second model, M was the dependent variable and diet/WHR intercept/slope + the independent variables (Path: X + covariates → M). If both the individual paths were significant, we ran the causal mediation analysis (Path: X → M → Y) using nonparametric bootstrapping to generate 95% confidence intervals (CIs) with 1,000 simulations.

**Appendix 8: Confounders**

All analyses were corrected for sex, MRI scanner model, age at the time of the MRI scan, years of education, BMI, mean arterial pressure, physical activity, and the Montreal Cognitive Assessment (MoCA) score, all measured at the MRI timepoint. Physical activity was measured using the Community Healthy Activities Model Program for Seniors (CHAMPS) questionnaire [14]. This self-reported questionnaire assesses the weekly frequency and duration of various activities, and the metabolic equivalent of task (met)/week of all activities was used as a confounder. Mean arterial pressure was derived as (systolic blood pressure+2*diastolic blood pressure)/3. RsfMRI analyses were additionally corrected for head motion and the voxel-wise grey matter density confounder. To quantify the quality instead of the quantity of food intake we included the total energy intake (kcal/day) as a confounder for the AHEI-2010 analyses.

**Appendix 9: Mean WM metrics and cognitive tests measurements**

***Table A3:*** *Mean outcome measures of extracted WM metrics and cognitive performance tests at MRI scan. Measures are specified as their mean ± their standard deviation. Diffusivity measures (MD, RD, AD) are multiplied with $10^3$.*

| | **AHEI-2010 sample included in functional connectivity analyses** | **AHEI-2010 sample included in WM connectivity analyses** | **WHR sample included in in functional connectivity analyses** | **WHR sample included in WM connectivity analyses** |
|---|---|---|---|---|
| N (% female) | 512 (21·29%) | 506 (21·54%) | 664 (19·88%) | 657 (20·09%) |
| Age (years) | 69·81 ± 5·11 | 69·78 ± 5·10 | 69·75 ± 5·07 | 69·72 ± 5·06 |

| | | | | |
|---|---|---|---|---|
| Working memory (verbal episodic memory - total recall) | 27·49 ± 4·68 | 27·52 ± 4·66 | 27·54 ± 4·60 | 27·57 ± 4·58 |
| Working memory (digit span total) | 30·63 ± 5·71 | 30·67 ± 5·71 | 30·79 ± 5·70 | 30·82 ± 5·69 |
| Semantic fluency | 22·49 ± 5·63 | 22·49 ± 5·64 | 22·34 ± 5·47 | 22·35 ± 5·47 |
| Lexical fluency | 15·75 ± 4·54 | 15·72 ± 4·55 | 15·77 ± 4·49 | 15·76 ± 4·49 |
| Executive function (digit coding) | 62·65 ± 13·29 | 62·74 ± 13·23 | 63·07 ± 13·21 | 63·14 ± 13·17 |
| Executive function (trail making) | 0·59 ± 0·48 | 0·58 ± 0·47 | 0·59 ± 0·45 | 0·58 ± 0·44 |
| WM global FA | | 0·48 ± 0·02 | | 0·29 ± 0·01 |
| WM global MD | | 0·68 ± 0·03 | | 0·36 ± 0·01 |
| WM global RD | | 0·49 ± 0·03 | | 0·28 ± 0·02 |
| WM global AD | | 1·08 ± 0·02 | | 0·64 ± 0·01 |
| WM ILF FA | | 0·14 ± 0·01 | | 0·14 ± 0·01 |
| WM cingulum FA | | 0·57 ± 0·03 | | 0·58 ± 0·03 |
| WM fornix FA | | 0·37 ± 0·09 | | 0·37 ± 0·09 |
| WM ILF MD | | 0·19 ± 0·01 | | 0·19 ± 0·01 |
| WM cingulum MD | | 0·63 ± 0·03 | | 0·63 ± 0·03 |
| WM fornix MD | | 1·38 ± 0·22 | | 1·43 ± 0·26 |
| WM ILF RD | | 0·13 ± 0·01 | | 0·13 ± 0·01 |
| WM cingulum RD | | 0·37 ± 0·03 | | 0·38 ± 0·03 |
| WM fornix RD | | 1·09 ± 0·27 | | 1·14 ± 0·31 |
| WM ILF AD | | 0·31 ± 0·01 | | 0·32 ± 0·01 |
| WM cingulum AD | | 1·14 ± 0·04 | | 1·15 ± 0·04 |
| WM fornix AD | | 1·96 ± 0·15 | | 2·00 ± 0·16 |

*Abbreviations: AHEI-2010 - Alternative-Health Eating Index 2010, WHR - waist-to-hip ratio, ILF - inferior longitudinal fasciculus, WM - white matter, FA - fractional anisotropy, MD - mean diffusivity, RD - radial diffusivity.*

### Appendix 10: Mean hippocampus connectivity maps

Across N=664 participants (i.e. participants of the WHR analyses), the right and left hippocampus showed strong correlations to each other, and to areas of the default mode network (DMN) such as the medial prefrontal cortex, posterior cingulate cortex/precuneus, and angular gyrus (see figure A2). We observed high hippocampal functional connectivity with most other cortical areas (r>0·3), whereas the functional connectivity to the brainstem and other subcortical structures was lower (r<0·2).

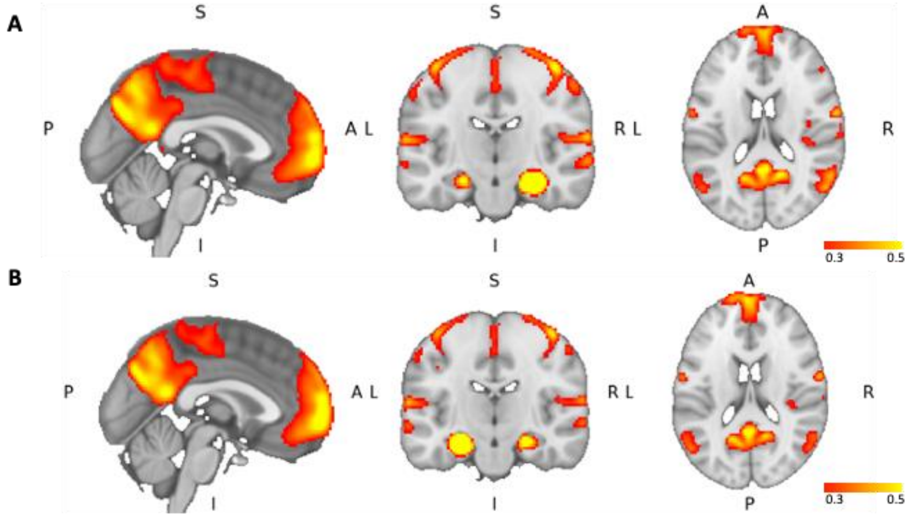

***Figure A2****: Mean functional connectivity from the right (A) and left (B) hippocampus mask to the rest of the brain. The intensity mask is thresholded to 0·3 for the lower limit and 0·5 for the upper limit. Maps are overlaid with the MNI-152 template. MNI coordinates are x=0, y=-18, z=18.*

### Appendix 11: Association of dietary quality and WHR with cognition

***Table A4**: Associations between the WHR (N=664), AHEI-2010 score (N=512), and cognitive outcomes. P-values are markers in bold for p<0·05 and include * p<0·0083 for the Bonferroni correction of multiple comparisons (six cognitive outcomes).*

| | Cognitive performance | β (SE) | CI | p-value |
|---|---|---|---|---|
| **intercept WHR** | Working memory (verbal episodic memory - total recall) | -0·0014 (0·0005) | -0·0024, -0·0004 | **0·0048*** |
| | Working memory (digit span total) | -0·0013 (0·0004) | -0·0021, -0·0006 | **0·00079*** |
| | Semantic fluency | -0·0010 (0·0004) | -0·0018, -0·0002 | **0·015** |
| | Lexical fluency | 0·0004 (0·0005) | -0·0006, 0·0013 | 0·43 |
| | Executive function (digit coding) | -0·0006 (0·0002) | -0·0009, -0·0002 | **0·00071*** |
| | Executive function (trail making) | 0·0052 (0·0050) | -0·0048, 0·0151 | 0·31 |
| **slope WHR** | Working memory (verbal episodic memory - total recall) | 0·0000 (0·0000) | -0·0001, 0·0001 | 0·68 |
| | Working memory (digit span total) | 0·0001 (0·0000) | 0·0000, 0·0002 | **0·023** |
| | Semantic fluency | -0·0000 (0·0000) | -0·0001, 0·0000 | 0·30 |
| | Lexical fluency | -0·0000 (0·0000) | -0·0001, 0·0000 | 0·33 |
| | Executive function (digit coding) | 0·0000 (0·0000) | -0·0000, 0·0001 | 0·16 |
| | Executive function (trail making) | 0·0005 (0·0005) | -0·0004, 0·0015 | 0·28 |
| **intercept AHEI-2010** | Working memory (verbal episodic memory - total recall) | 0·0244 (0·0701) | -0·1134, 0·1621 | 0·73 |
| | Working memory (digit span total) | 0·0247 (0·0578) | -0·0890, 0·1383 | 0·67 |
| | Semantic fluency | 0·0067 (0·0582) | -0·1077, 0·1211 | 0·91 |
| | Lexical fluency | -0·0892 (0·0702) | -0·2271, 0·0488 | 0·20 |
| | Executive function (digit coding) | 0·0038 (0·0249) | -0·0451, 0·0527 | 0·88 |
| | Executive function (trail making) | -0·9812 (0·6853) | -2·3276, 0·3653 | 0·15 |
| **slope AHEI-2010** | Working memory (verbal episodic memory - total recall) | 0·0030 (0·0032) | -0·0033, 0·0094 | 0·35 |
| | Working memory (digit span total) | 0·0036 (0·0027) | -0·0016, 0·0088 | 0·18 |
| | Semantic fluency | 0·0003 (0·0027) | -0·0050, 0·0056 | 0·90 |
| | Lexical fluency | 0·0073 (0·0032) | 0·0010, 0·0137 | **0·024** |
| | Executive function (digit coding) | 0·0001 (0·0011) | -0·0022, 0·0024 | 0·94 |
| | Executive function (trail making) | 0·0095 (0·0317) | -0·0528, 0·0718 | 0·76 |

*Abbreviations: CI - confidence interval.*

### Appendix 12: Mediation analyses for all WM brain outcomes

***Table A5:** Results of 95 % confidence intervals of mediation (1000x Bootstrapping). All models are adjusted for confounders (see Methods) and include complete participants information. Significant mediation effects are indicated in bold and with * for p<0·0071 (Bonferroni correction for multiple comparisons across seven mediators). Tested mediators were: global (MD and RD), ILF (FA, MD, and RD), and Cingulum (MD and RD)). We tested the indirect effect when the linear regression of the individual paths (X → M, and M → y) showed significant predictions (see appendix 8).*

| Y (cognition) | M (WM brain outcomes) | X → M (CI) | M → Y (CI) | Indirect effect (ACME: X → M → Y) | Direct / Total effect (X → Y) | Proportion mediated |
|---|---|---|---|---|---|---|
| verbal episodic memory | global RD | 564·19 (259·52, 868·86) p=0·0003 | 0 (-0·0, -0·0) p=0·0362 | -0·0001 (-0·0003, 0·0) p=0·048 | -0·0014 (-0·0025, 0·0) p=0·010 | 0·1019 (-0·0035, 0·34) p=0·058 |
| digit span | global MD | 571·02 (213·50, 928·54) p=0·0018 | 0 (-0·0, -0·0) p=0·0011 | -0·0002 (-0·0003, 0·0) p<2e-16 | -0·0013 (-0·0022, 0·0) p=0·002 | 0·1205, (0·0298, 0·36) p=0·002 |
| | global RD | 534·93 (228·11, 841·75) p=0·0007 | 0 (-0·0, -0·0) p=0·0003 | -0·0002 (-0·0004, 0·0) p<2e-16 | -0·0013 (-0·0022, 0·0) p<2e-16 | **0·1464, (0·0411, 0·38) p<2e-16*** |
| | FA in ILF | -1·12 (-1·82, -0·42) p=0·0017 | 0·0002 (0·0001, 0·0002) p=0·0007 | -0·0002 (-0·0003, 0·0) p=0·008 | -0·0013 (-0·0021, 0·0) p<2e-16 | 0·1254, (0·0242, 0·34) p=0·008 |
| | MD in ILF | 0·74 (0·22, 1·25) p=0·005 | -0·0002 (-0·0003, -0·0) p=0·0069 | -0·0001 (-0·0003, 0·0) p=0·006 | -0·0013 (-0·0021, 0·0) p=0·002 | 0·0899, (0·0094, 0·26) p=0·008 |
| | RD in ILF | 0·81 (0·3278, 1·29) p=0·001 | -0·0002 (-0·0003, -0·0001) p=0·0016 | -0·0002 (-0·0004, 0·0) p=0·004 | -0·0013 (-0·0021, 0·0) p=0·004 | 0·1228, (0·0267, 0·39) p=0·008 |
| | MD in Cingulum | 0·26 (0·06, 0·45) p=0·0096 | -0·0004 (-0·0007, -0·0001) p=0·0227 | -90·42e-05 (-20·43e-04, 0·0) p=0·038 | -10·35e-03 (-0·0021, 0·0) p<2e-16 | 0·0698, (0·0029, 0·25) p=0·038 |
| | RD in Cingulum | 0·21 (0·04, 0·37) p=0·015 | -0·0005 (-0·0009, -0·0002) p=0·0055 | -0·0001 (-0·000254, 0·0) p=0·03 | -0·0013 (-0·0021, 0·00) p<2e-16 | 0·0800, (0·0061, 0·26) p=0·03 |

| | | | | | | |
|---|---|---|---|---|---|---|
| digit coding | global MD | 609·97 (254·24, 965·70) p=0·0008 | 0 (-0·0, -0·0) p=0·0028 | -6·88e-05 (-1·43e-04, 0·0) p=0·006 | -5·68e-04 (-9·01e-04, 0·0) p=0·004 | 0·1210, (0·0260, 0·34) p=0·010 |
| | global RD | 569·77 (264·38, 875·16) p=0·0003 | 0 (-0·0, -0·0) p=0·0006 | -8·63e-05 (-1·62e-04, 0·0) p<2e-16 | -5·68e-04 (-8·91e-04, 0·0) p<2e-16 | **0·1520, (0·0466, 0·43) p<2e-16*** |
| | FA in ILF | -1·20 (-1·90, -0·51) p=·0007 | 0·0001 (0·0, 0·0001) p=0·0082 | -6·15e-05 (-1·31e-04, 0·0) p=0·008 | -5·68e-04 (-9·19e-04, 0·0) p=0·002 | 0·1080, (0·0191, 0·31) p=0·010 |
| | RD in ILF | 0·87 (0·39, 1·35) p=0·0004 | -0·0001 (-0·0001, -0·0) p=0·0125 | -6·09e-05 (-1·28e-04, 0·0) p=0·022 | -5·68e-04 (-9·22e-04, 0·0) p=0·002 | 0·1070, (0·0130, 0·31) p=0·024 |
| | MD in Cingulum | 0·27 (0·08, 0·47) p=0·006 | -0·0002 (-0·0003, -0·0) p=0·0161 | -4·55e-05 (-1·09e-04, 0·0) p=0·028 | -5·68e-04 (-9·43e-04, 0·0) p=0·002 | 0·0801, (0·0024, 0·28) p=0·030 |
| | RD in Cingulum | 0·21 (0·04, 0·38) p=0·0133 | -0·0003 (-0·0005, -0·0002) p=0·0001 | -6·57e-05 (-1·40e-04, 0·0) p=0·026 | -5·68e-04 (-9·27e-04, 0·0) p<2e-16 | 0·1160, (0·0118, 0·35) p=0·026 |

*Abbreviations: WM - white matter, FA - fractional anisotropy, MD - mean diffusivity, RD - radial diffusivity, X - predictor variable, y - outcome variable, M - mediator variable, ILF - inferior longitudinal fasciculus, CI - confidence interval, ACME - average causal mediation effect.*
